# Similar does not mean same: ERP correlates of processing mental and physical experiencer verbs in Malayalam complex constructions

**DOI:** 10.1101/2025.05.21.655344

**Authors:** S. Shalu, R. Muralikrishnan, Kamal Kumar Choudhary

## Abstract

The present study examined the neurophysiological correlates of processing mental experiencer verbs and physical experiencer verbs in Malayalam complex constructions, in which the subject argument assumed the experiencer role. Event-related brain potentials (ERPs) were recorded as twenty-eight first-language speakers of Malayalam read intransitive sentences with the two types of experiencer verbs. The subject case either matched the requirements of the verb in the critical stimuli, thereby rendering the sentence acceptable, or it violated the requirements of the verb, and thus rendering the sentence unacceptable. Both mental and physical experiencer verbs engendered negativity effects in the 400–600 ms time-window when the subject case did not match the verb’s requirements. Additionally, mental experiencer verbs evoked a LAN-like effect in the same time window regardless of grammaticality. Thus, even though both kinds of experiencer verbs are processed qualitatively similarly, inherent differences between mental and physical experiencer verbs in Malayalam persist and are discernible.

## 1. Introduction

Interest in the processing of different verb types and their cross-linguistic applicability has been growing over the past few decades. While early studies focused on broad classifications of verbs, more recent research has explored increasingly detailed subcategories, highlighting the nuanced distinctions in verb processing. One area of investigation has been the processing of subcategories of concrete verbs (Bedny et al., 2008; Kemmerer et al., 2008; Thomas et al., 2012; Dalla Volta, 2014; Repetto et al., 2015), and their comparison with abstract verbs^1^ (Binder et al., 2005, Goldberg et al., 2007, Hoffman et al., 2015, Hoffman et al., 2010, Perani et al., 1999, Sabsevitz et al., 2005), though relatively few studies have examined the processing of different types of abstract verbs (Rodríguez-Ferreiro et al. 2011; Muraki et al., 2020. Dryer et al., 2015; Dryer et al., 2018). A notable contribution in this regard is Muraki et al. (2020), who argued that treating abstract verbs homogeneously limits our understanding of their underlying representation.

Most of the existing ERP studies on abstract verbs were conducted within the framework of embodied cognition theories. A central finding of this line of research is that different types of abstract verbs are represented differently in the brain (Muraki et al., 2020; Rodríguez-Ferreiro et al. 2011; Dryer et al., 2015; Dryer et al., 2018). However, there is limited research on processing different categories of abstract verbs based on their argument structure and realization. One of the verb categories examined in this regard is experiencer verbs and their major subcategories such as subject experiencer verbs (SE) and object experiencer verbs (OE). Several behavioral studies have examined the processing differences between SE and OE, as well as how these verbs differ from action verbs, revealing that SE are easier to process than OE, because of their less complicated alignment encoding grammatical relations and thematic roles (Linda Cupples, 2000; Kretzschmar et al., 2012; Gattei et al., 2015; Gattei et al., 2018; Gattei et al., 2022; Mateu 2022; Wilson & Dillon, 2022). No study to date has examined the processing of experiencer verbs within the SE category, except for Shalu et al. (2025), who investigated mental experiencer verbs (ME, indicating mental experiences like happiness, sadness, etc.) and physical experiencer verbs (PE, indicating physical experiences like hunger, thirst, cold, etc.) in Malayalam, a South Dravidian language, which clearly distinguishes between these verbs in its syntax.

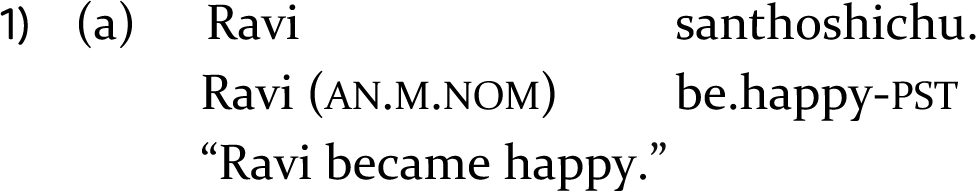
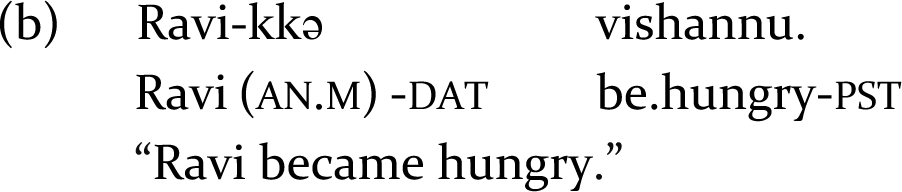

In simple experiencer constructions, ME require a nominative subject (1a), while PE require a dative subject (1b) in Malayalam. Shalu et al. (2025) crossed subject case and verb type in their design, and reported N400s for both ME and PE when the subject was anomalous. However, they also found differences in peak latency of the effect between ME and PE, as well as an influence of sentence-final acceptability on the ERPs for the non-anomalous conditions, suggesting subtle but robust differences in processing ME versus PE.

However, a key limitation of their design is, it is not possible to disentangle the effect of the verb type per se from the effect of differing expectations due to the sentence-initial nominative versus dative subject interacting with verb type. This is because the subject case (say, nominative) that constituted a violation for PE was, by contrast, the non-anomalous subject case for ME, and vice versa. A potential way to address the issue would be to examine ME and PE in complex predicates, since both verb types require a dative subject in these constructions (2a–b).

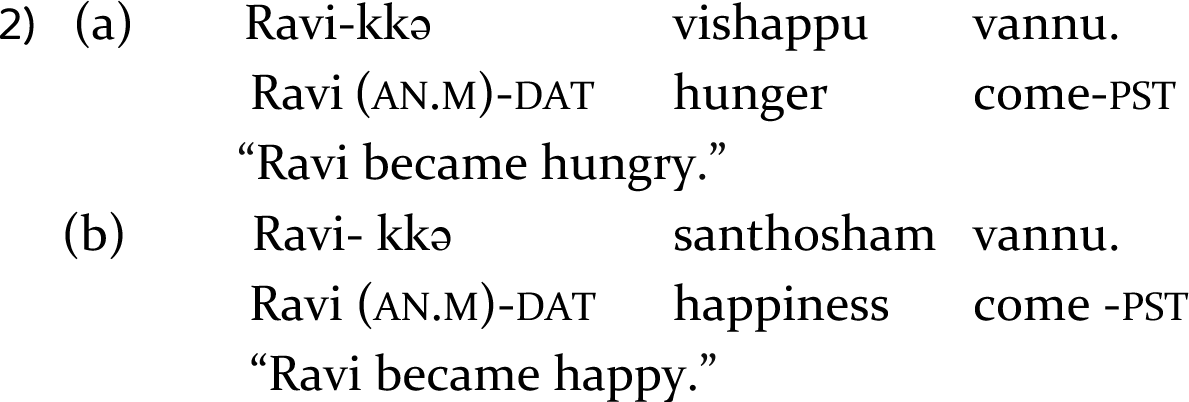

### 1.1. The present study

We employed complex intransitive constructions in the present study, formed using an event nominal (physical: ‘*pani*’ feve*r,* or mental: ‘*santhosham*’ happiness) that provides most of the meaning and activates its argument structure, followed by a semantically underspecified light verb (‘*vannu*’ come). We employed a 2 x 2 factorial design similar to Shalu et al. (2025), manipulating the subject case (nominative or dative) and predicate type (ME or PE), keeping the light verb identical across conditions. Since the sentence-initial subject was uniformly dative in all non-anomalous conditions, and nominative in all violation conditions, any ERP differences at the verb for ME violations versus PE violations would be directly attributable to a processing difference between the two verb types.

Based on the theoretical perspective on ME and PE in Malayalam as well as results from Shalu et al. (2025), we hypothesize that if ME and PE in Malayalam complex constructions are processed similarly, then qualitatively similar ERP correlates should ensue for both verb types. This would be in line with the theoretical literature on experiencer verbs, as the argument structure is identical for both ME and PE (SubjectDAT-EXP- VerbMENT/PHY). Following Shalu et al. (2025), we expected an N400 effect for both the violation conditions compared to their respective correct counterparts as a marker of the violation of an interpretively relevant linguistic rule. Alternatively, since a large part of the predicative meaning originates from the event nominal within the construction (Butt, 2010), and the nominals in the study represent two very different types of experiences, i.e., mental vs. physical (Jayaseelan, 2004), we should observe qualitative neurophysiological differences between the two verb types. Whilst the exact nature of this difference remains to be seen, observing such a difference would add to the claim from Shalu et al. (2025) that the two types of experiencers show inherent processing differences despite their overall processing similarity in Malayalam. Further, if the differences in peak latency for ME versus PE in Shalu et al. (2025) were due to inherent distinctions between these verb types per se, we should expect to see similar differences in the present study. However, if these differences were stemming from nominative vs dative subject case interacting with verb type, no such difference should ensue.

## 2. Method

### 2.1. Participants

Twenty-eight first-language speakers of Malayalam (mean age = 27.8; 11 female and 17 male) participated in the experiment in exchange for payment. Data from 7 participants were excluded from analysis due to excessive artifacts (see supplementary material for further details).

### 2.2. Materials

We employed critical stimuli in 4 conditions (Table 1), with 36 sentences in each critical condition, resulting in a total of 144 critical sentences. These were interspersed with fillers and pseudorandomized for presentation. The supplementary material provides further details.

**Table 1.** A sample set of experimental stimuli. Experiencer light verbs are indicated as Mental_LV (mental experiencer) and Physical_LV (physical experiencer) and case markers are indicated as Nom (nominative) and Dat (dative). Dat-Mental_LV and Dat-Physical_LV are grammatically correct sentences, Nom-Mental_LV and Nom-Physical_LV are ungrammatical.

| Condition | Sample Stimulus |
| --- | --- |
| Dat-Mental_LV | Ravi-kkə santhosham vannu.<br>Ravi (AN.M)-DAT happiness come-PST<br>“Ravi became happy.” |
| Nom-Mental_LV | *Ravi santhosham vannu.<br>Ravi (AN.M.NOM) happiness come-PST<br>“Ravi became happy.” |
| Dat-Physical_LV | Ravi-kkə vishappu vannu.<br>Ravi (AN.M)-DAT hunger come- PST<br>“Ravi became hungry.” |
| Nom-Physical_LV | *Ravi vishappu vannu.<br>Ravi (AN.M.NOM) hunger come-PST<br>“Ravi became hungry.” |

### 2.3. Procedure

Participants were seated in a dimly lit, sound attenuated room during the experiment, which included a short practice. Stimuli were presented using E-prime 2.0 (Psychology Software Tools, Pittsburgh, PA) in a rapid serial visual presentation. Participants performed an acceptability judgement and a probe task after each trial. The supplementary material provides further details.

### 2.4. EEG recording, Pre-processing and Statistical processing

Scalp activity was recorded by means of 32 Ag/AgCI electrodes fixed at the scalp using Hydrocel geodesic Sensor Net 32, with Cz as online reference. EEG data was pre- processed using EEGLAB (Version 14; Delorme & Makeig, 2004, sccn.ucsd.edu) in MATLAB (Version R2023b; The MathWorks, Inc.). Data epochs were extracted at the critical position (verb; -200 to 1200 ms) from the continuous recordings for each participant and analyzed using the eeguana package (Version 0.1.11.9001; Nicenboim, 2018) in R (Version 4.4.3; R Core Team, 2024).

ERP Data Analysis: The mean amplitudes in the time-window of interest were statistically analyzed using the single trial EEG epochs at the verb for each critical condition by fitting linear mixed effects models in R (Version 4.4.3, R Core Team 2024) using the lme4 package (Bates et al., 2015). The statistical models included the fixed factors Case (Nominative vs Dative) and Verb type (Mental experiencer vs Physical experiencer), as well as the topographical factor Regions of Interest (ROI). We included the mean amplitude from the 200 ms pre-stimulus period (-200 to 0 ms) as a (scaled and centred) covariate in the model for each data epoch (Alday, 2019). Categorical fixed factors used sum contrasts (scaled sum contrasts for 2-level factors), so that the coefficients represent deviations from the grand mean (Schad et al., 2020). The ROIs were defined by clustering topographically adjacent electrodes in 6 lateral and 2 midline regions. The lateral ROIs were as follows: Left-Frontal, which included electrodes E3 and E11 (which, in the 10-20 electrode system, would have been equivalent to F3 and F7); Left-Central, which included electrodes E5 and E13 (C3 and T7); Left-Parietal, which included electrodes E7 and E15 (P3 and P7); Right-Frontal, which included electrodes E4 and E12 (F4 and F8); Right-Central, which included electrodes E6 and E14 (C4 and T8); and Right-Parietal, which included electrodes E8 and E16 (P4 and P8). Mid-Fronto-Central, which included E17 and E28 (Fz and ∼FCz); and Mid-Parieto-Occipital, which included E19, E20, E9, and E10 (Pz, Oz, O1 and O2), were the midline ROIs. The supplementary material provides further details.

## 3. Results

### 3.1. Behavioral data

The mean acceptability ratings and probe detection accuracy for the critical conditions were calculated for the trials included in the ERP analysis. As Table 2 shows, the non-anomalous conditions were rated as highly acceptable (> 98 %; ceiling), whereas the acceptability for the violation conditions was very low across the board (< 21 %). The probe detection accuracy reached ceiling for all the conditions. Figure 1 shows raincloud plots (Allen, Poggiali, Whitaker, Marshall, & Kievit, 2021) of the behavioural acceptability judgements.

**Figure 1.**
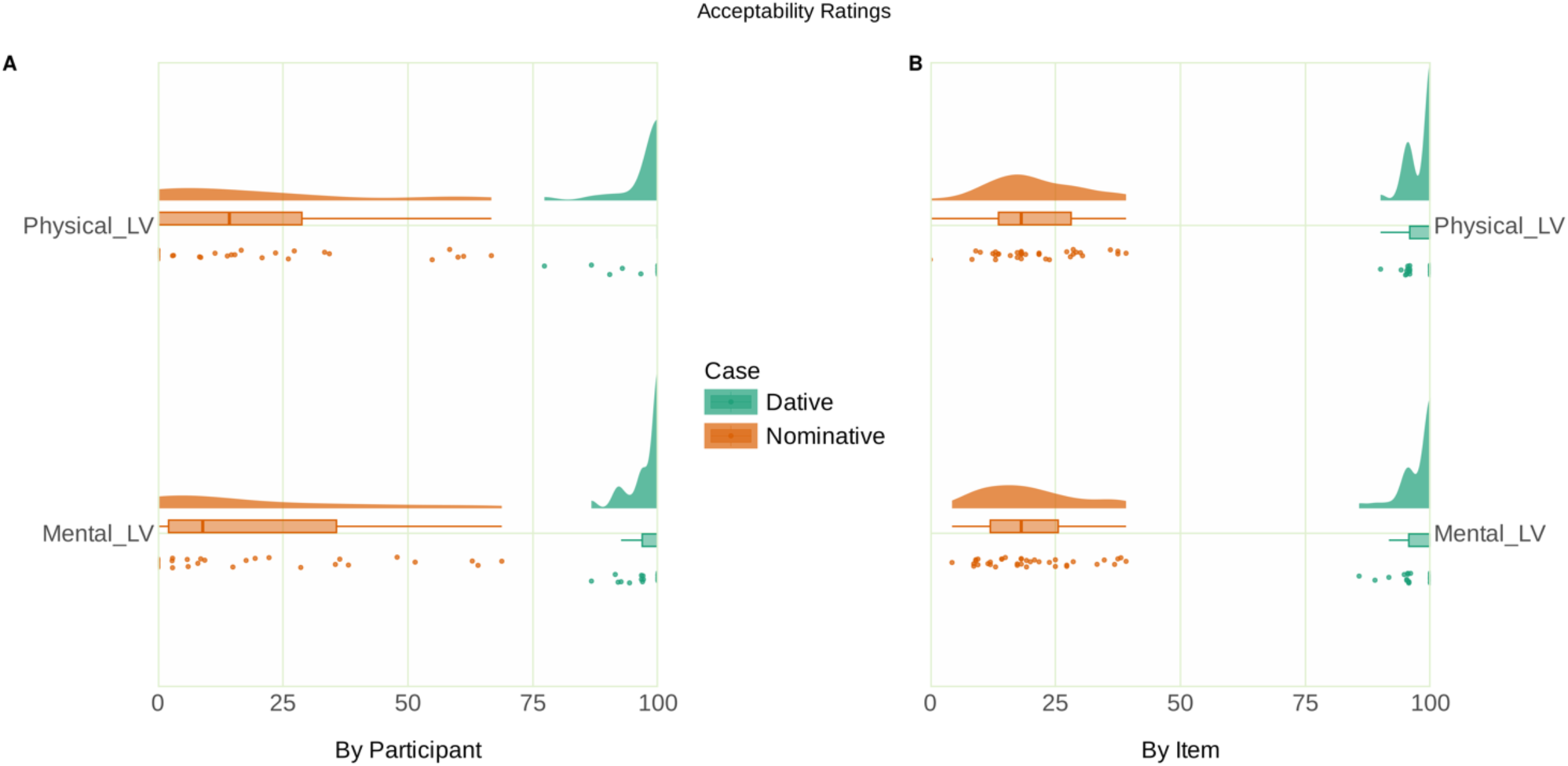
**Raincloud plot of the acceptability ratings. Panel A shows the by-participant variability of acceptability ratings, with the individual data points representing the mean by-participant acceptability of each case and verb type combination. Panel B shows the by-item variability of acceptability ratings, with the individual data points representing the mean by-item acceptability of each case and verb type combination.**

**Table 2.**
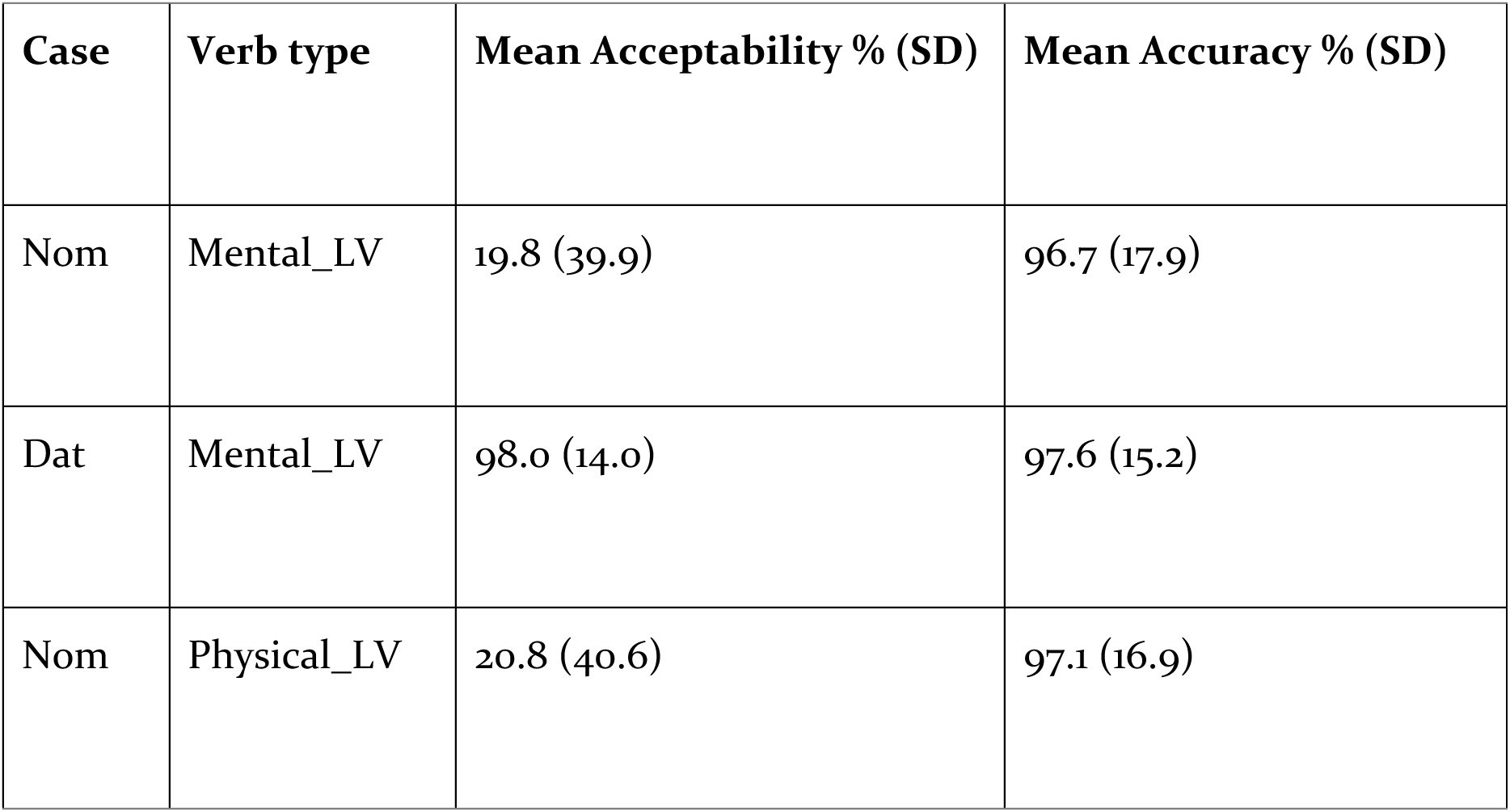
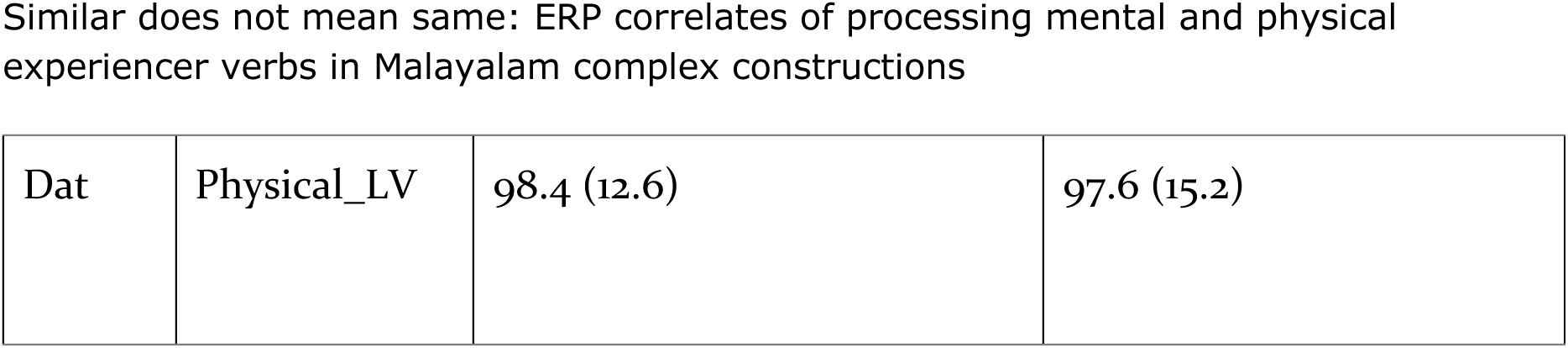
Mean acceptability ratings and probe detection accuracy.

The behavioral acceptability and accuracy were analyzed by fitting generalized linear mixed models using the lme4 package in R. Categorical fixed factors used scaled sum contrasts (effect coding). In the analysis of acceptability data, the statistical model included the fixed factors Case (Nominative vs Dative) and Verb type (Mental experiencer vs Physical experiencer), with random intercepts for participants and items, and by-participant random slopes for the effect of Case, Verb type and their interaction term. Type II Wald chi-square tests of the fitted model (AIC = 1571.012) of the acceptability data showed a main effect of Case (χ^2^(1) = 109.66, p < 0.001, s = 82.83) and the interaction of Verb x Case (χ^2^(1) = 7.49, p= 0.006, s = 7.33). Estimated marginal means on the response scale were computed on the model using the emmeans package (Lenth, 2021) to resolve this interaction, which showed that, for both verb types, the estimates for the conditions with a dative subject were higher compared to those for conditions with a nominative subject. Pairwise contrasts of these estimates within each verb type revealed a simple effect of Case for both mental experiencer light verbs (estimate **=** 0.878, SE **=** 0.038, p < 0.001, s **=** 373.79) and physical experiencer light verbs (estimate **=** 0.873, SE **=** 0.040, p **<** 0.001, s **=** 346.06). These findings reflect the substantial difference in acceptability between conditions with dative subjects, which engendered very high acceptability, versus conditions with nominative subjects, which were rated as hardly acceptable.

In the analysis of the probe detection accuracy, models with an interaction term of the fixed factors Verb and Case, as well as those with random slopes were singular. The model with Verb and Case as fixed factors with by-participant and by-item random intercepts (AIC = 816.1534) did not detect any effect involving Verb or Case (s < 2.5), reflecting the highly similar (ceiling) accuracy of probe detection across conditions.

### 3.2. ERP data

The ERPs at the verb are shown in Fig. 2. Visual inspection of the ERP data indicated a negativity effect in the violation conditions relative to the non-anomalous ones. Based on both the ERP components identified in previous research (Shalu et al., 2025) and visual inspection, we selected the 400–600 ms time-window for analysis. The single trial ERP mean amplitudes extracted in the analysis time-window from a total of 3238 data epochs entered the analysis. The raw data collected during the experiment, the pre-processing pipeline used, and the pre-processed data are available in the data repository online.

**Figure 2.**
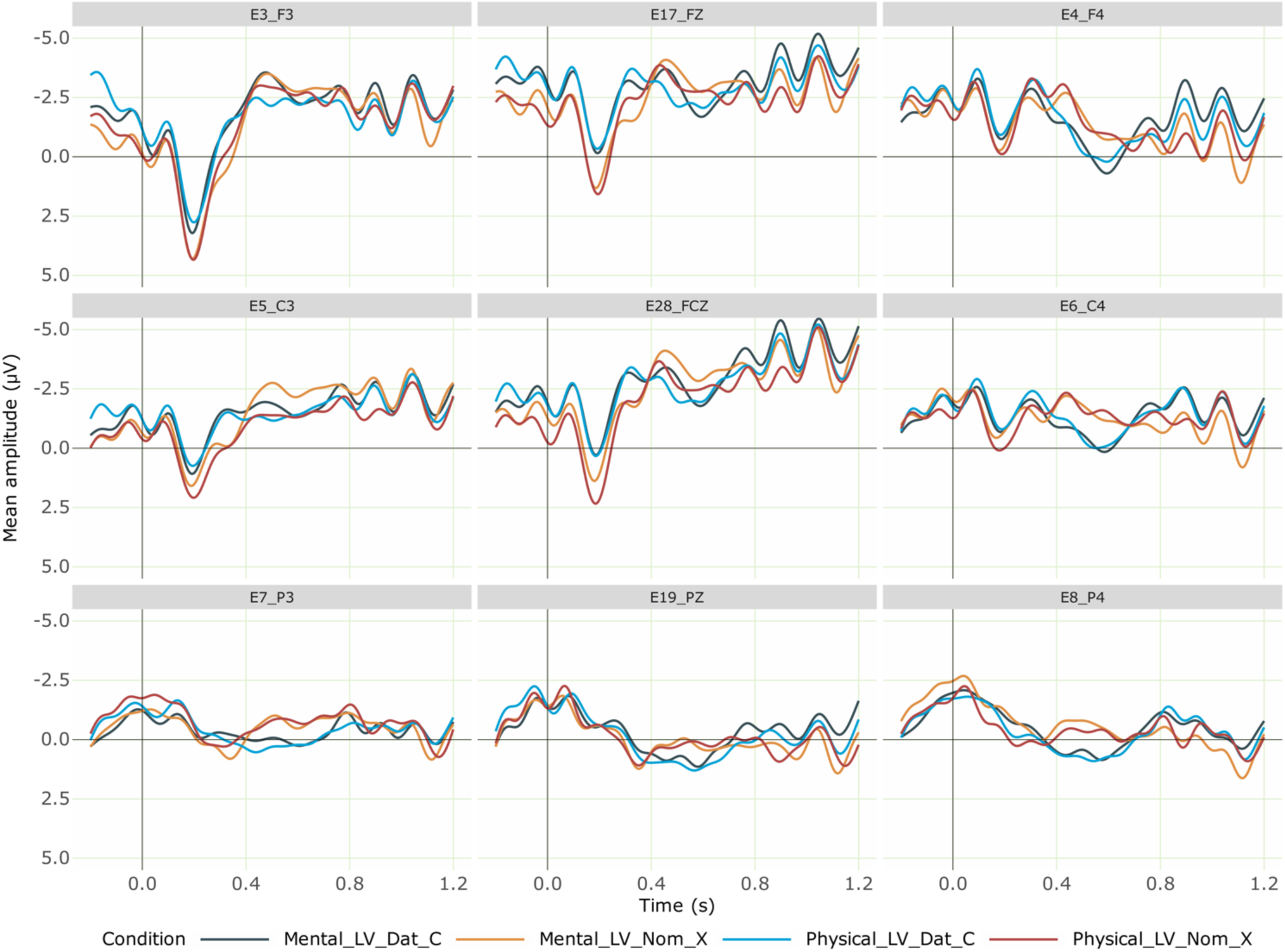
Grand averaged ERPs at the verb for the critical conditions from 28 participants. Negativity is plotted upwards; the time axis runs from -0.2 s to **1.2** s (i.e., -200 ms to 1200 ms) with 0 being the onset of the critical verb. The dark blue line represents the correct mental experiencer light verb (with a dative subject), while the orange line shows the incorrect mental experiencer light verb (with a nominative subject), which elicited a centro-parietal negativity effect. The light blue line indicates the correct physical experiencer verb (with a dative subject) and the red line represents the incorrect physical experiencer verb (with a nominative subject), which elicited a centro-parietal negativity effect. Regardless of grammaticality, conditions with mental experiencer light verbs elicited a further negativity effect in the left anterior region compared to physical experiencer light verbs.

We computed a linear mixed-effects model with the fixed factors Verb type, Case of the subject noun, ROI, the -200-0 ms pre-stimulus baseline mean amplitude as a covariate (scaled and centred), and by-participant and by-item random intercepts. The analysis code and full model outputs are available as R notebooks in the analysis repository online. Type II Wald Chi-squared tests on this model (AIC = 434325.1) showed a main effect of Case (χ^2^(1) = 6.84, p = 0.008, s = 6.81) and interaction effects of ROI x Verb (χ^2^(1) = 14.79, p = 0.03, s = 4.69) and ROI x Case (χ^2^(1) = 14.82, p = 0.03, s = 4.70). Estimated marginal means on the response scale were computed on the model using the emmeans package (Lenth, 2021) to resolve these interactions. The pairwise contrasts of estimates for Case within each level of ROI revealed simple effects of Case in the right-frontal (estimate = 1.245, SE = 0.364, p < 0.001, s = 10.60), right-central (estimate = 1.067, SE = 0.364, p = 0.003, s = 8.19), right-parietal (estimate = 1.012, SE = 0.364, p = 0.005, s = 7.50), mid-fronto-central (estimate = 0.947, SE = 0.365, p = 0.009, s = 6.71) and mid-parieto-occipital (estimate = 0.566, SE = 0.319, p = o.07, s = 3.71) regions. The estimates for nominative subjects were more negative than those for dative subjects in these regions. The pairwise contrasts of estimates for Verb type within each level of ROI revealed a simple effect of Verb type in the left-frontal region (estimate = -0.839, SE= 0.364, p = 0.02, s = 5.54). The estimate for mental experiencer verb was more negative than that for physical experiencer verb in this region. A more complex model in which the by-participant random slopes for Case, Verb type, and their interaction term were included in the random effects specification^2^ showed that this pattern of results remained largely intact despite numerical differences. In sum, both mental and physical experiencer verbs elicited a negativity effect when the subject was nominative as opposed to dative. Further, a general verb-type effect ensued for mental experiencer verbs, which elicited a LAN-like effect compared to physical experiencer verbs, regardless of grammaticality.

## 4. Discussion

We have presented an ERP experiment on Malayalam, which aimed to examine whether the complex ME and PE are processed qualitatively similarly. The behavioural task demonstrated a clear link between grammaticality and acceptability, with violation conditions leading to a drastically low acceptability compared to grammatical sentences. The ERP results showed a negativity effect in the 400–600 ms time-window, which can be plausibly interpreted as an instance of an N400 effect for violations involving both the verb types. More interestingly however, there was an unexpected left anterior negativity (LAN-like) effect in the same time-window for ME regardless of grammaticality; that is, the effect was a general verb type effect observed for both anomalous and non-anomalous ME in comparison to PE. In addition to this contrast to findings from Shalu et al. (2025), the current study did not observe any peak latency differences in the negativity effect between ME and PE. This can be attributed to the fact that both conditions begin with a nominative argument, thus generating identical expectations about possible continuations. And in both cases, these expectations are violated in a similar manner when the parser encounters the ME or PE verb. This finding supports the notion that the latency differences reported by Shalu et al. (2025) may have been attributable to the violation of expectations based on the nominative versus dative subject arguments, rather than reflecting an inherent distinction between ME and PE per se.

### 4.1. N400 effect

The N400 effect found for both the experiencer verbs are consistent with Shalu et al. (2025). As in their study, we interpret the negativity effects that we found in the present study as a consequence of a violation of interpretively relevant rules, in line with Choudhary et al. (2009) and Nieuwland et al. (2013).

In Malayalam, a clear morphosyntactic constraint governs the case marking of the subject argument of ME and PE in complex constructions. Regardless of whether the experience meant by the experiencer verb is mental or physical, it is expressed as though it is moving from an abstract space to the subject argument, which serves as the experiencer-goal of the abstract movement. In this respect, the dative subject experiencer is similar to other dative arguments that are the targets/goals of some kind of movement (see sentence 5 -7). Consequently, the subject is in dative case, which is the general obligatory marker for targets of movement in Malayalam (Nizar, 2010; Mohanan, 1990; Krishnan, 2019; Jayaseelan, 2004). The violation of this interpretively relevant rule in the present study led to an N400 effect for both verb types, regardless of the experience conveyed by the verb.

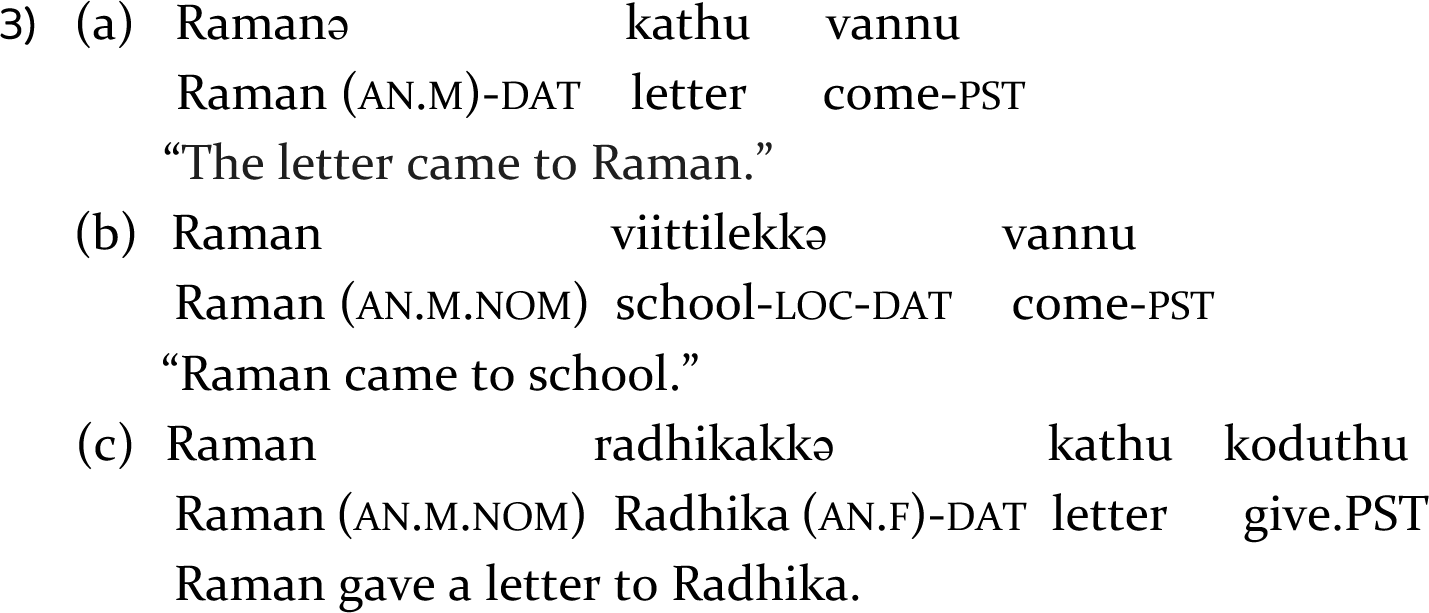

### 4.2. LAN-like effect

The classic LAN effect is typically elicited by anomalous inflectional morphology (Gunter et al., 1997). It is generally understood as an indicator of the detection of morpho-syntactic inconsistencies (Münte et al., 1997; Friederici, 2002, see Molinaro et al., 2011 for a review), but has also been found for syntactic violations (Friederici et al., (1996); Hagoort et al., (2003); Hahne & Friederici, (1999); Münte et al.,(1993), well-formed ambiguous constructions (Hagoort and Brown, 2000), and syntactically complex constructions (Kluender and Kutas,1993; Münte et al., 1998). The LAN is typically observed in the same time-window as the N400, with often the only difference between the two effects being the topography.

It is however unclear if we can interpret the LAN-like effect in our study as an instance of the LAN. Crucially for present purposes, previous studies on abstract verb processing have nevertheless shown that it activates left-lateralized brain regions (Rodriguez-Ferriero et al., 2011; Binder et al., 2005; Sabsevitz et al., 2005; Noppeney & Price, 2004; Perani, Schnur, et al., 1999). In line with this, it is plausible that the LAN-like effect observed in our study for ME can be explained by the degree of abstractness and non-bodily involvement associated with these verbs compared to PE. ME refer to internal states, often linked to emotions, thoughts etc., which are intangible and cannot be directly observed. By contrast, PE describe physical sensations or states.

Compared to ME which do not directly engage with the body, PE are closely linked to the physical body and the tangible sensations it can perceive and feel. That is, even though PE are abstract, they have some observable effects in the body that make them more perceivable compared to relatively more abstract mental experiences.

For instance, an individual suffering from fever may have a flushed face and sweating, an individual who is experiencing hunger may have a growling sound from the stomach, etc. Thus, the LAN-Like effect observed for ME can be plausibly attributed to their more abstract nature compared to PE.

Alternatively, the LAN-like effect for ME in comparison to PE can be explained in terms of the ± control/agency associated with both the verb types. ME refer to a state where the experiencer has volitional control/agency over their mental state, allowing them to change or influence it without taking physical action. For instance, someone can willfully suppress sadness to maintain composure. In contrast, PE describe a state where the experiencer has little control/agency over the situation and must take tangible actions to change it. For example, a person experiencing freezing cold must actively do something to alleviate the discomfort, such as wearing warm clothes or putting the heater on. In light of this, it is plausible to consider the LAN-like effect observed in our study as a marker of the control/agency difference between ME and PE. However, this interpretation remains speculative, and more studies comparing agentive verbs with non-agentive verbs would be needed to gain further insights in this regard.

In conclusion, the present study explored the processing of mental and physical experiencer verbs in Malayalam complex experiencer constructions. Results revealed a negativity effect for subject case violations involving mental as well as physical experiencer verbs, suggesting that they are processed qualitatively similarly. However, we also found a LAN-like effect for mental experiencer verbs regardless of grammaticality compared to physical experiencer verbs. In sum, although both mental and physical experiencer verbs are processed qualitatively similarly, our data nevertheless provides converging evidence to conclude that effects of inherent differences between mental and physical experiencer verbs in Malayalam are still discernible even in syntactically identical constructions involving the two verb types.

## Data availability

Data repository. The raw data collected during the experiment, the pre-processing pipeline used, and the pre-processed data are available here: https://zenodo.org/records/14986233?token=eyJhbGciOiJIUzUxMiJ9.eyJpZCI6ImViODI2Y2IzLWM5MTUtNDc5OC1iNTBlLTQ0ZGVjYWI4M2U2YyIsImRhdGEiOnt9LCJyYW5kb20iOiIyYmJhMmY4ZmRjYmE4MjAxNjA2OWQxYjJmNWU4YzIzOCJ9.bXrtqZEiasDf2kd788hblBXp3E4DDJk8Ef0Qq0NK5n6bVO8SC_kWoEILQgj4mpdbFmIO8h5FP8UtGn6bXNXzgA

Analysis repository. The analysis code and full model outputs of all the analyses reported are available as R notebooks here: https://zenodo.org/records/15478572?token=eyJhbGciOiJIUzUxMiJ9.eyJpZCI6IjFiZDEyZTlhLWM2YTUtNGU1NC05NDQ1LWExYjBlNTU4YjhhMyIsImRhdGEiOnt9LCJyYW5kb20iOiJiMjIwYWM2ZDMyMGIzOTU3MDJhOTg0YzdkM2ZmNzhlMyJ9.Kpg1TkJP-ljNqLQPd8U6DM_bvcA-hq33T3XGdcqre1HXAPXpd4WJGrXYe8Lpx4oMsMRLCAKigmTZUTRH5VnBCQ

## Ethics Statement

The research protocol for this experiment was approved by the Institute Ethics Committee (IEC) at the Indian Institute of Technology Ropar, where the study took place. Participants provided informed written consent upon arrival at the lab. All procedures conducted during the experiment adhered to the guidelines and regulations set forth by the committee.

## Author Contributions

SS: Conceptualization, Investigation, Methodology, Project administration, Resources, Software, Data curation, Formal analysis, Validation, Visualization, Writing – original draft, Writing – review & editing.

RM: Conceptualization, Software, Data curation, Formal analysis, Funding acquisition, Supervision, Validation, Visualization, Writing – review & editing.

KKC: Conceptualization, Methodology, Funding acquisition, Supervision, Validation, Visualization, Writing – review & editing.

## Acknowledgements

The work was supported by the Department of Science and Technology, Govt. of India, through a project to Dr. Kamal Kumar Choudhary (SR/CSI/141/2012). We would like to acknowledge the Indian Institute of Technology Ropar (IIT Ropar) for the research facility maintained through an ISIRD grant to Dr Kamal Kumar Choudhary [IITRPR/Acad./2216]. We acknowledge the Max Planck Society for covering the Article Processing Costs that is provided through an institutional arrangement available for submitting and corresponding authors affiliated with the Max Planck Society.

## Supplementary Material

### A1. Participants

A total of 28 first-language speakers of Malayalam (mean age = 27.8; age range = 18–40; 11 female and 17 male), primarily students and staff at the Indian Institute of Technology Ropar, residing in Ropar, India, participated in the experiment in exchange for payment. Prior to the experiment, participants were informed about the study, and written consent was obtained for the use of their data for academic purposes. The research protocol was approved by the Ethics Committee (Human) of the Indian Institute of Technology Ropar. All participants were right-handed, as determined by an adapted version of the Edinburgh Handedness Inventory (Oldfield, 1971) in Malayalam. They had normal or corrected-to-normal vision and were free from known neurological disorders at the time of the experiment. All participants were native Malayalam speakers, having acquired the language before the age of six. Additionally, most spoke other languages. Data from 7 participants were excluded from analysis due to excessive artifacts.

### A2. Materials

In the critical conditions, we employed 9 mental experiencer (ME) complex predicates and 9 physical experiencer (PE) complex predicates. Each verb was repeated 4 times with 4 different nouns (2 masculine and 2 feminine nouns) which resulted in 36 sets of sentences with 144 critical sentences as in Table 2 in the main article. The light verb was identical across all sentences (“vannu-come”). The sentence-initial noun was either nominative (null case marking) or dative marked. Since Malayalam dative case marker has two allomorphs (“-kkə” or the “-ə” markers^3^), we used an equal number of nouns requiring either the “-kkə” or the “-ə” marker (18 each). In these 144 critical sentences, 72 sentences were grammatically correct (dative good), and 72 sentences were grammatically incorrect (nominative bad). Additionally, we used 288 fillers, which were constructed to introduce diverse structures in the stimuli so as to avoid strategic responses from participants. They also served to counterbalance the number of grammatical and ungrammatical sentences with different subject types (72 good nominative sentences, 72 bad nominative sentences, 72 bad dative sentences and 72 good dative sentences). The fillers were interspersed with the critical stimuli and pseudorandomized for presentation during the experiment.

### A3. Procedure

The experiment began with a practice session, followed by the actual experiment, with all activities, including electrode preparation and stimulus presentation, taking approximately 3 hours. Participants were first briefed about the procedure and the tasks they would perform and were provided with a printed instruction sheet.

However, the specific research question was not disclosed to ensure unbiased data collection. Participants then filled out a consent form to provide informed consent for their participation, along with a Malayalam version of the Edinburgh Handedness Inventory to assess their handedness. Next, their head measurements were taken, and the Hydrocel GSN net was applied to their scalp. They were seated comfortably in a soundproof chamber, positioned 1 meter from a 20” LCD monitor where the stimuli were displayed. Stimuli presentation was managed using E-prime 2.0 (Psychology Software Tools, Pittsburgh, PA).

The structure of a trial in the experiment was as follows: sentences were presented visually, chunk-by-chunk, at the centre of the monitor screen. At the start of each trial, participants saw a "+" fixation sign at the centre of the screen for 1000 ms. This was followed by a blank screen for 100 ms. Each chunk was presented for 650 ms, followed by an interstimulus interval (ISI) of 100 ms (Muralikrishnan & Idrissi, 2021; Demiral et al., 2008; Bornkessel-Schlesewsky et al., 2020). After the fixation point, a noun phrase (NP) appeared on the screen, followed by a verb (critical position for the study). After this, participants performed two tasks. Since a violation paradigm was used in the study, they first completed an acceptability judgment task. A “???” sign appeared on the screen to signal the acceptability judgment task. They had to press the green button on the response pad if they found the preceding sentence acceptable, or the red button if they found it unacceptable. Additionally, to ensure the participants were paying attention to the sentence, they also completed a probe task. After finishing the acceptability judgment task or once 1500 ms had passed, a probe word appeared on the screen for 2000 ms. Participants were required to press the green button if the probe word was present in the preceding sentence or the red button if it was not. For instance, an experimental sentence like “menuvinu (thanuppu vannu)” (“Meenu became cold”) might be followed by a probe word “meenuvinu” or an incorrect probe word like “kappal” (“ship”). Half of the probe words required a ‘yes’ response, while the other half required a ‘no’ response.

The experiment was conducted in two versions: in one version, the correct response was mapped to the left key and the incorrect response to the right key, while in the other version, this mapping was reversed. Participants were instructed to avoid blinking during the stimulus presentation but were allowed to blink while performing the tasks. Prior to the actual experiment, they underwent a practice session to familiarize themselves with the trial structure, although none of the real experimental stimuli were included in the practice session. The experimental session was divided into 9 blocks, each containing 48 sentences, with short breaks at the end of each block. At the conclusion of the experiment, participants were asked to complete a questionnaire about their experience.

#### A.3.1. EEG recording, Pre-processing and Statistical processing

Scalp activity was recorded by means of 32 Ag/AgCI electrode fixed at the scalp by means of Hydrocel geodesic Sensor Net 32 channel (reference). Cz served as the online reference. To monitor EOG (Electro-oculogram) data, electrodes were placed as follows: for horizontal eye movements, electrodes were positioned at the outer canthus of each eye, and for vertical eye movements, electrodes were placed below the eyes. The interelectrode impedance was maintained below 50 kΩ (with an amplifier input impedance greater than 1 GOhm) (Ferree et al., 2001). All EEG and EOG data were amplified using a Net Amps 400 Amplifier and recorded at a sampling rate of 500 Hz.

The EEGLAB toolbox (Version 14; Delorme & Makeig, 2004, sccn.ucsd.edu) in MATLAB (Version R2023b; The MathWorks, Inc.) was used for the pre-processing of EEG data. The data was down-sampled to 250 Hz and filtered with a 0.3–20 Hz band- pass filter to eliminate slow drifts. These settings encompass the typical 0.5–5 Hz range for language-related ERP activity (Delorme, 2023; Roehm et al., 2002) and have been widely used in cross-linguistic ERP studies on language processing. Channels on the outer extent of the face and head were eliminated after this data was offline- referenced to the mean of the two mastoids. A copy of this original data that had been high-pass filtered at 1 Hz was subjected to an Independent Component Analysis (ICA, Iriarte et al., 2003).

Before sending this data to an ICA computation using the extended Infomax algorithm, bad channels were removed. The ICLabel plugin (Pion-Tonachini, Kreutz- Delgado & Makeig, 2019) was then used to test the resultant Independent Components (ICs) in order to identify and mark artefactual ICs. The bad channels that were removed prior to computing the ICA were interpolated from the remaining channels. The weights calculated during ICA were then copied to the original data. The ICs identified as artefactual were then removed from the original data. The eeguana package (Version 0.1.11.9001; Nicenboim, 2018) was then used to import the data into R (Version 4.4.3; R Core Team, 2024) for statistical analysis and epoching.

Data epochs were extracted from the continuous recordings for each participant, focusing on the critical position of the verb. These epochs spanned from 200 ms before the onset to 1200 ms after the onset (i.e., from -200 to 1200 ms). Epochs were discarded if the amplitude exceeded a threshold of 100 μV in either direction, or if the difference between the minimum and maximum amplitudes within a 200 ms window surpassed the 100 μV threshold. Additionally, trials in which the acceptability judgment task was not completed were excluded. Data from participants with too few remaining trials were also removed from further analysis. After these rejections, there were approximately 28 to 29 valid trials per condition per participant, so the number of trials included in the analysis was similar across conditions. In total, 3238 data epochs/trials were included in the analysis across participants, with a median of 807 trials per critical condition. For visualization, the valid epochs were averaged across items within each condition for each participant, and then grand averages were calculated across participants. These grand averages were smoothed with an 8 Hz low- pass filter to generate the ERP plots for the NP and verb in each condition.

ERP Data Analysis: The mean amplitudes in the time-window of interest were statistically analysed using the single trial EEG epochs at the NP and the verb for each critical condition. This was done by fitting linear mixed effects models in R (Version 4.4.3, R Core Team 2024) using the lme4 package (Bates et al., 2015). The statistical models included the fixed factors Case (Nominative vs Dative) and Verb type (Mental experiencer vs Physical experiencer), as well as the topographical factor Regions of Interest (ROI). The ROIs were defined by clustering topographically adjacent electrodes in 6 lateral and 2 midline regions. The lateral ROIs were as follows: Left- Frontal, which included electrodes E3 and E11 (which, in the 10-20 electrode system, would have been equivalent to F3 and F7); Left-Central, which included electrodes E5 and E13 (C3 and T7); Left-Parietal, which included electrodes E7 and E15 (P3 and P7); Right-Frontal, which included electrodes E4 and E12 (F4 and F8); Right-Central, which included electrodes E6 and E14 (C4 and T8); and Right-Parietal, which included electrodes E8 and E16 (P4 and P8). Mid-Fronto-Central, which included E17 and E28 (Fz and ∼FCz); and Mid-Parieto-Occipital, which included E19, E20, E9, and E10 (Pz, Oz, O1 and O2), were the midline ROIs.

Instead of using a traditional subtraction-based baseline correction, we included the mean amplitude from the 200 ms pre-stimulus period (-200 to 0 ms) as a covariate (after scaling and centering) in the model for each data epoch. This approach was taken to account for and remove potential baseline differences in the statistical analysis, as suggested by Alday (2019). However, we did not interpret any effects involving pre-stimulus amplitudes, consistent with the fact that these were not part of our hypotheses. This also aligns with Alday’s (2019) recommendation that "additional covariates can be included as controls without further interpretation". For the contrasts related to categorical factors, we used sum contrasts (scaled sum contrasts for 2-level factors), so that the coefficients represent deviations from the grand mean (Schad et al., 2020). Following modern statistical recommendations, we do not use the term ‘statistically significant’ or its variants for describing effects based on p-value thresholds (Wasserstein et al., 2019), but report precise p-values as continuous quantities (e.g., p = 0.06 rather than p < 0.08), unless a value is “below the limit of numerical accuracy of the data”, in which case, we report it as p < 0.001 (Amrhein et al., 2019, p.266). Further, we supplement the p-values by transforming them into s- values (Shannon information, surprisal, or binary logworth) and report s = – log2(p), which provides on an absolute scale, a nonprobability measure of the information provided by a p-value (Shannon, 1948; Greenland, 2019). In other words, “the s-value provides a gauge of the information supplied by a statistical test” and has the advantage of providing “a direct quantification of information without” requiring prior distributions as input (Rafi & Greenland, 2020, p.6).

### A4. ERPs at the sentence-initial subject noun

Figure. S1 shows the ERPs at the position of the sentence-initial subject noun. We computed a linear mixed effects model with fixed factors Case of the subject noun, ROI, the -200-0 ms pre-stimulus baseline mean amplitude as a covariate (scaled and centred), and by-participant and by-item random intercepts. The analysis code and full model outputs are available as R notebooks in the analysis repository online. Type II Wald Chi-squared tests on this model (AIC = 399983.75) showed an interaction effect of ROI x Case (χ^2^(1) = 35.56, p = 0.00001, s = 16.80). Estimated marginal means on the response scale were computed on the model using the emmeans package (Lenth, 2021) to resolve this interaction. The pairwise contrasts of estimates for Case within each level of ROI revealed simple effects of Case in the left-frontal region (estimate = - 0.817, SE = 0.326, p = 0.01), right-parietal region (estimate = 0.613, SE = 0.326, p = 0.06) and mid-fronto-central (estimate = -0.731, SE = 0.326, p = 0.02) region. The estimate for the dative subjects were more negative than those for the nominative subjects in the left-frontal and mid-fronto-central region. In the right-parietal region, the estimate for dative subjects is less positive than that for the nominative subjects. As the effects at the sentence-initial noun do not form part of our original hypotheses at the verb position, we refrain from interpreting them further. A more complex linear mixed-effects model was computed, including by-participant random slopes for Case^4^, which showed that this pattern of results remained largely intact despite numerical differences.

**Figure S1.**
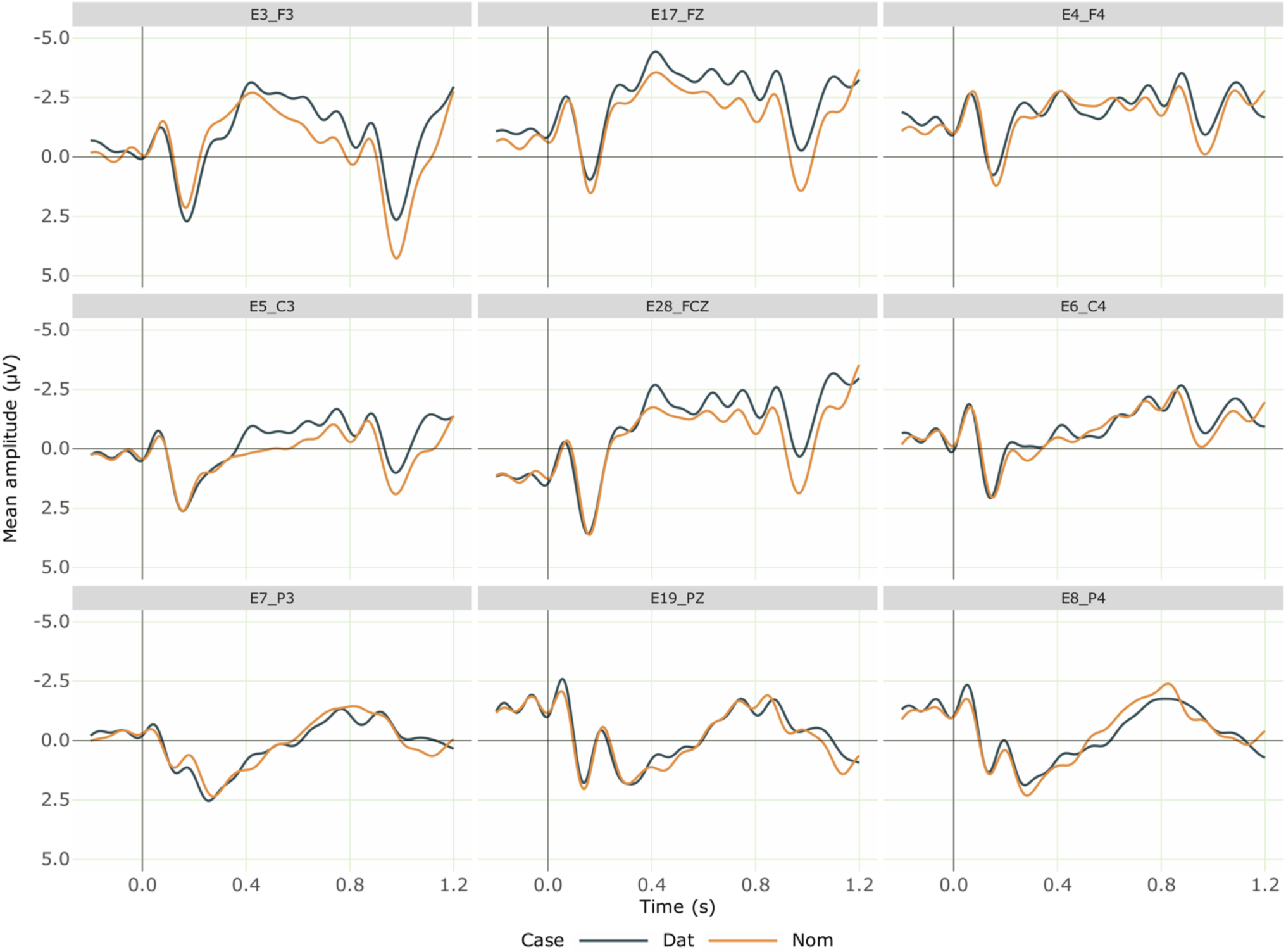
Grand averaged ERPs at the nominative and dative sentence-initial subject nouns from 28 participants. Negativity is plotted upwards; the time axis runs from -0.2 s to 1.2 s (i.e., -200 ms to 1200 ms) with 0 being the onset of the noun. The dark blue line shows the ERPs for the dative nouns and the orange line shows that for the nominative nouns.

### A5. Absence of late positivity effects at the Verb

Figure. S2 and Figure. S3 show the ERP at the verb for mental and physical experiencer verb conditions separately respectively. The study revealed no late positivity effects, which is a similar finding to that reported by Shalu et al. (2025). We observed only an N400 despite the understanding that the ill-formed sentences typically elicit a P600 effect (Bornkessel-Schlesewsky & Schlesewsky, 2008). The absence of P600 effects can be potentially explained by the resolvability of the violation condition (Frenzel et al., 2010; Bornkessel et al., 2011). The violation constructions in the experiment can be non-anomalous when we use them in a different context such as a serial verb construction, as in (A-C).

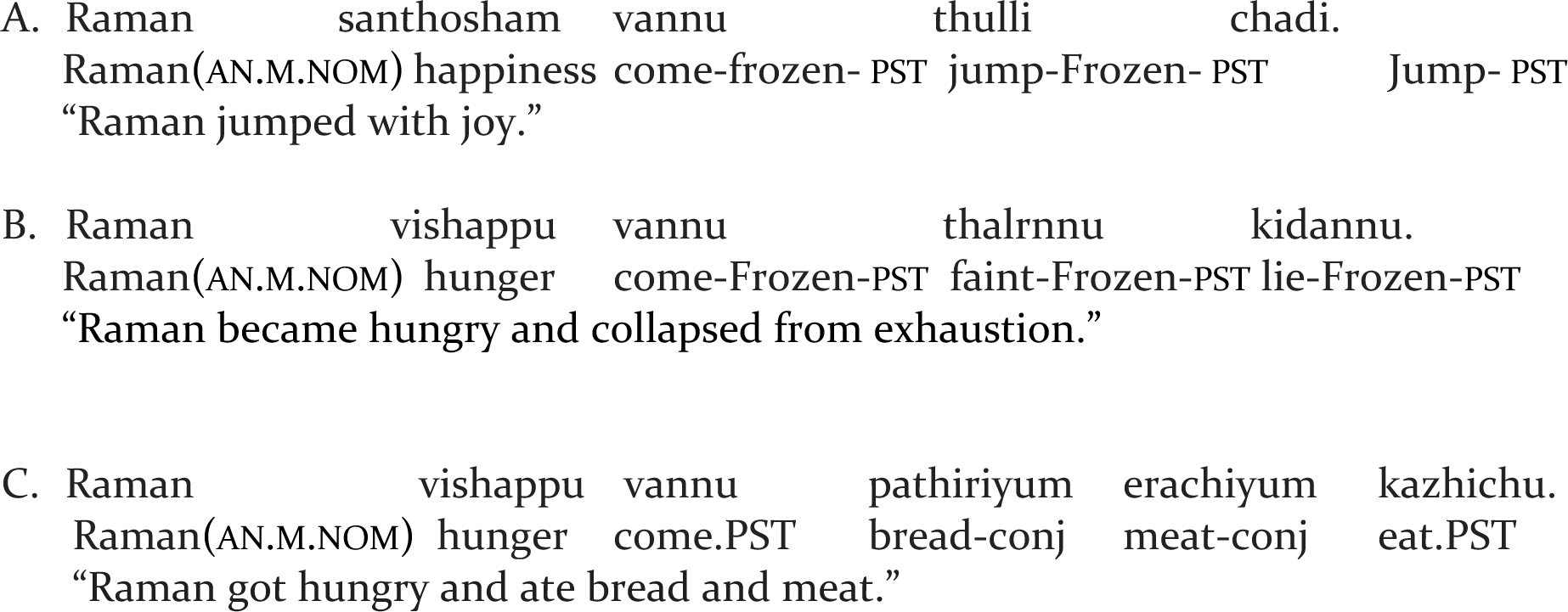

#### ERPs at the verb: Mental experiencer verbs

**Figure S2.**
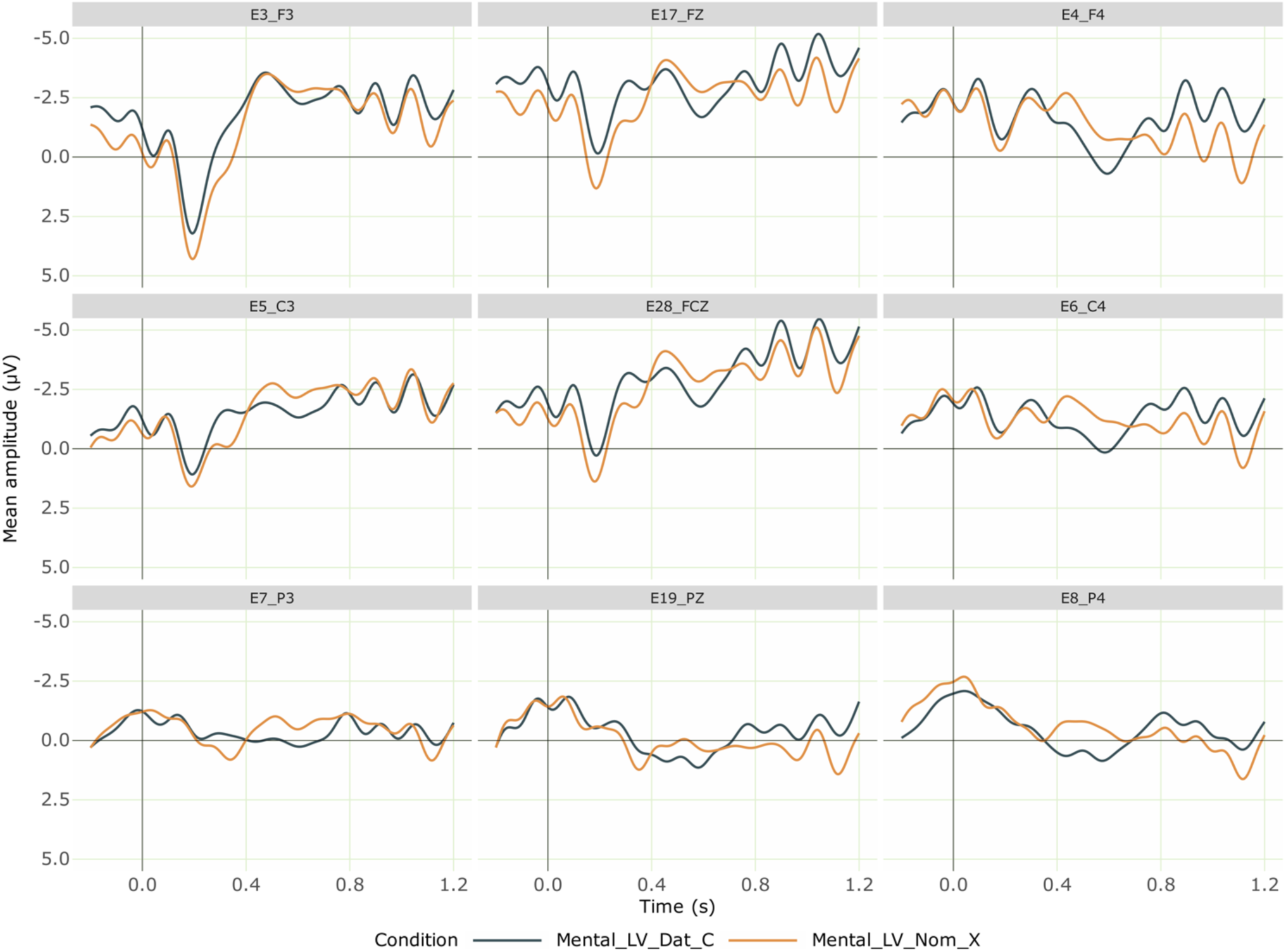
Grand averaged ERPs at the position of Mental experiencer light verb comparing Dative mental experiencer light verb (Correct) and Nominative mental experiencer light verb (Incorrect) from 28 participants. Negativity is plotted upwards; the time axis runs from -0.2 s to 1.2 s (i.e., -200 ms to 1200 ms) with 0 being the onset of the critical verb. The dark blue line shows the ERPs for the correct Mental experiencer verb (with a dative subject) and the orange line shows that for incorrect mental experiencer verb (with a nominative subject), which elicited a negativity effect.

#### ERPs at the verb: Physical experiencer verbs

**Figure S3.**
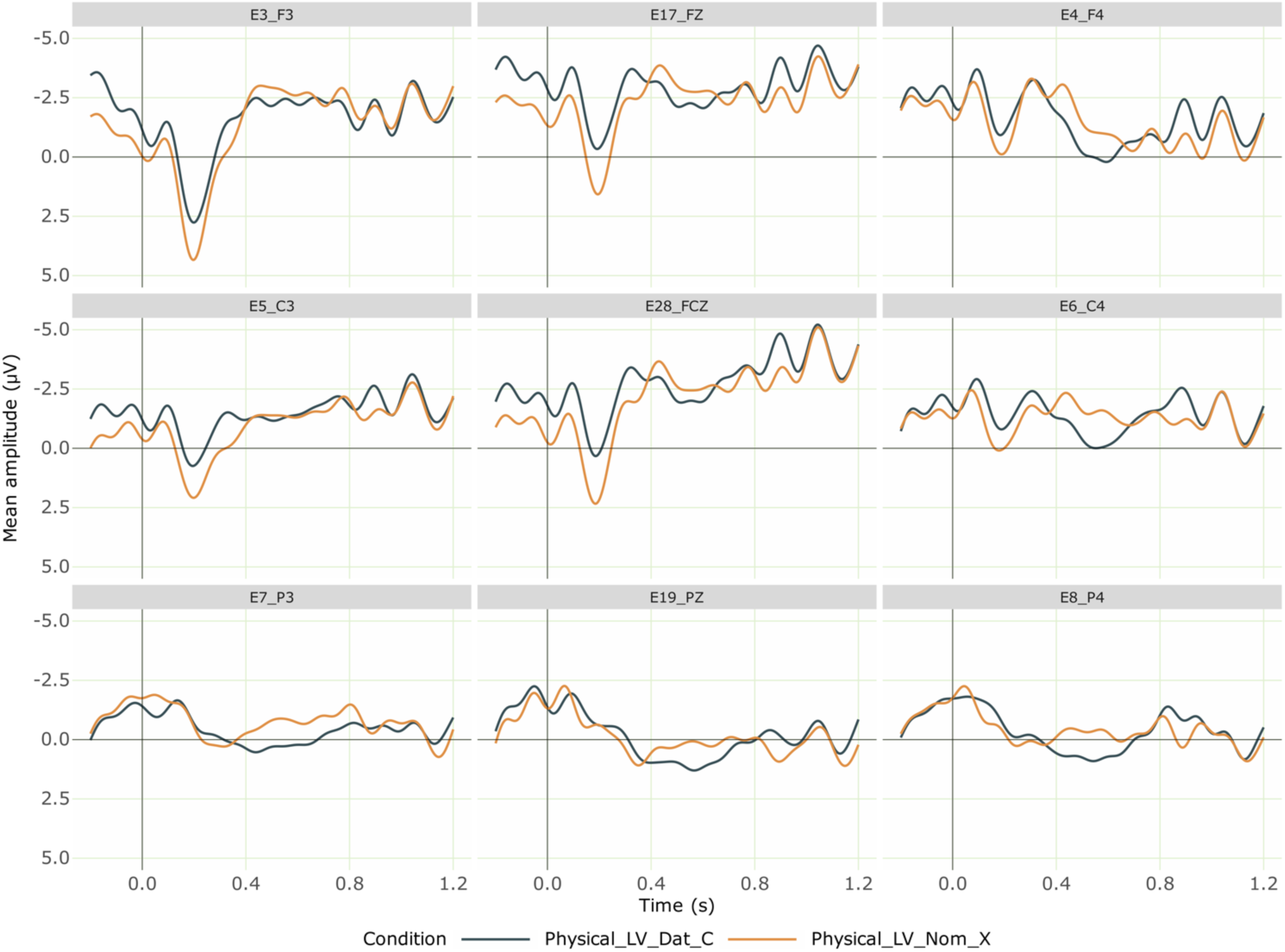
Grand averaged ERPs at the position of Physical experiencer light verb comparing Dative Physical experiencer light verb (Correct) and Nominative Physical experiencer light verb (Incorrect) from 28 participants. Negativity is plotted upwards; the time axis runs from -0.2 s to 1.2 s (i.e., -200 ms to 1200 ms) with 0 being the onset of the critical verb. The dark blue line shows the ERPs for the correct Physical experiencer light verb (with a dative subject) and the orange line shows that for incorrect Physical experiencer light verb (with a nominative subject), which elicited a negativity.

#### ERPs at the verb: All conditions

**Figure S4.**
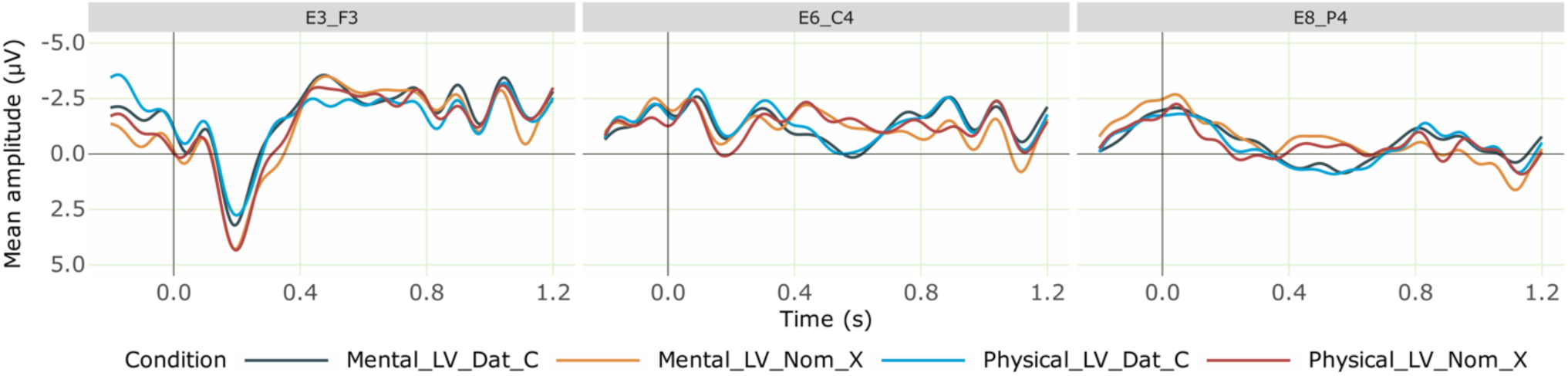
Grand averaged ERPs at the verb for the critical conditions from 28 participants. Negativity is plotted upwards; the time axis runs from -0.2 s to 1.2 s (i.e., -200 ms to 1200 ms) with 0 being the onset of the critical verb. The dark blue line represents the correct mental experiencer light verb (with a dative subject), while the orange line shows the incorrect mental experiencer light verb (with a nominative subject), which elicited a centro-parietal negativity effect. The light blue line indicates the correct physical experiencer verb (with a dative subject) and the red line represents the incorrect physical experiencer verb (with a nominative subject), which elicited a centro-parietal negativity effect. Regardless of grammaticality, conditions with mental experiencer light verbs elicited a further negativity effect in the left anterior region compared to physical experiencer light verbs.

## Footnotes

1 We consider verbs that do not refer to tangible motor actions as abstract verbs and assume abstractness to be a matter of degree.

2 Since the more complex model was nearly equivalent to the simpler model in terms of fit, predictive accuracy, and the amount of variance explained, we retain the simpler model for further interpretation. The full model output for the complex model is provided for reference in the analysis repository online.

3 The usage of these case suffixes is purely based on the phonology (last syllable) of the noun to which these are attached, regardless of the gender, number and other properties of the noun concerned. The allomorph “ə” is used for nouns or their derivation ending in “n”, while “-kkə” is applied elsewhere (Rajendran,1977).

4 The full model output for the complex model is provided for reference in the analysis repository online.

## Reference

1. Alday, P. M. (2019). How much baseline correction do we need in ERP research? Extended GLM model can replace baseline correction while lifting its limits. In Psychophysiology (Vol. 56, Issue 12). Wiley. 10.1111/psyp.13451

2. Allen, M., Poggiali, D., Whitaker, K., Marshall, T. R., van Langen, J., & Kievit, R. A. (2021). Raincloud plots: a multi-platform tool for robust data visualization. In Wellcome Open Research (Vol. 4, p. 63). F1000 Research Ltd. 10.12688/wellcomeopenres.15191.2

3. Amrhein, V., Trafimow, D., & Greenland, S. (2019). Inferential Statistics as Descriptive Statistics: There Is No Replication Crisis if We Don’t Expect Replication. In The American Statistician (Vol. 73, Issue sup1, pp. 262–270). Informa UK Limited. 10.1080/00031305.2018.1543137

4. Barber, H., & Carreiras, M. (2005). Grammatical gender and number agreement in Spanish: An ERP comparison. Journal of cognitive neuroscience, 17(1), 137–153.

5. Bates, D., Mächler, M., Bolker, B., & Walker, S. (2015). Fitting Linear Mixed-Effects Models Usinglme4. In Journal of Statistical Software (Vol. 67, Issue 1). Foundation for Open Access Statistic. 10.18637/jss.v067.i01

6. Bedny, M., Caramazza, A., Grossman, E., Pascual-Leone, A., & Saxe, R. (2008). Concepts are more than percepts: the case of action verbs. Journal of Neuroscience, 28(44), 11347–11353.

7. Binder, J. R., Westbury, C. F., McKiernan, K. A., Possing, E. T., & Medler, D. A. (2005). Distinct brain systems for processing concrete and abstract concepts. Journal of cognitive neuroscience, 17(6), 905–917.

8. Bornkessel-Schlesewsky, I., Roehm, D., Mailhammer, R., & Schlesewsky, M. (2020). Language Processing as a Precursor to Language Change: Evidence From Icelandic. In Frontiers in Psychology (Vol. 10). Frontiers Media SA. 10.3389/fpsyg.2019.03013

9. Bornkessel, I., McElree, B., Schlesewsky, M., & Friederici, A. D. (2004). Multi-dimensional contributions to garden path strength: Dissociating phrase structure from case marking. Journal of Memory and Language, 51(4), 495–522.

10. Bornkessel, I., Schlesewsky, M., & Friederici, A. D. (2002). Beyond syntax: Language-related positivities reflect the revision of hierarchies. NeuroReport, 13(3), 361–364.

11. Bornkessel, I., Schlesewsky, M., & Friederici, A. D. (2003). Eliciting thematic reanalysis effects: The role of syntax-independent information during parsing. Language and Cognitive Processes, 18(3), 269–298.

12. Butt, M. (2010). The light verb jungle: Still hacking away. Complex predicates: Cross-linguistic perspectives on event structure, 48-78.

13. Choudhary, K. K., Schlesewsky, M., Roehm, D., & Bornkessel-Schlesewsky, I. (2009). The N400 as a correlate of interpretively relevant linguistic rules: Evidence from Hindi. In Neuropsychologia (Vol. 47, Issue 13, pp. 3012–3022). Elsevier BV. 10.1016/j.neuropsychologia.2009.05.009

14. Cupples, L. (2002). The structural characteristics and on-line comprehension of experiencer-verb sentences. In Language and Cognitive Processes (Vol. 17, Issue 2, pp. 125–162). Informa UK Limited. 10.1080/01690960143000001

15. Dalla Volta, R., Fabbri-Destro, M., Gentilucci, M., & Avanzini, P. (2014). Spatiotemporal dynamics during processing of abstract and concrete verbs: An ERP study. Neuropsychologia, 61, 163–174.

16. Delorme, A. (2023). EEG is better left alone. In Scientific Reports (Vol. 13, Issue 1). Springer Science and Business Media LLC. 10.1038/s41598-023-27528-0

17. Delorme, A., & Makeig, S. (2004). EEGLAB: an open source toolbox for analysis of single-trial EEG dynamics including independent component analysis. Journal of neuroscience methods, 134(1), 9–21.

18. Demiral, Ş. B., Schlesewsky, M., & Bornkessel-Schlesewsky, I. (2008). On the universality of language comprehension strategies: Evidence from Turkish. In Cognition (Vol. 106, Issue 1, pp. 484–500). Elsevier BV. 10.1016/j.cognition.2007.01.008

19. Gattei, C. A., Alvarez, F., París, L., Wainselboim, A., Sevilla, Y., & Shalom, D. (2022). Why bother? What our eyes tell about psych verb (non) causative constructions. In Glossa Psycholinguistics (Vol. 1, Issue 1). California Digital Library (CDL). 10.5070/g601138

20. Gattei, C. A., Dickey, M. W., Wainselboim, A. J., & París, L. (2015). The thematic hierarchy in sentence comprehension: A study on the interaction between verb class and word order in Spanish. In Quarterly Journal of Experimental Psychology (Vol. 68, Issue 10, pp. 1981–2007). Sage Publications. 10.1080/17470218.2014.1000345

22. Gattei, C. A., Sevilla, Y., Tabullo, Á. J., Wainselboim, A. J., París, L. A., & Shalom, D. E. (2017). Prominence in Spanish sentence comprehension: an eye-tracking study. In Language, Cognition and Neuroscience (Vol. 33, Issue 5, pp. 587–607). Informa UK Limited. 10.1080/23273798.2017.1397278

23. Gattei, C. A., Tabullo, Á., París, L., & Wainselboim, A. J. (2015). The role of prominence in Spanish sentence comprehension: An ERP study. In Brain and Language (Vol. 150, pp. 22–35). Elsevier BV. 10.1016/j.bandl.2015.08.001

24. Gunter, T. C., Friederici, A. D., & Schriefers, H. (2000). Syntactic gender and semantic expectancy: ERPs reveal early autonomy and late interaction. Journal of cognitive neuroscience, 12(4), 556–568.

25. Hagoort, P., & Brown, C. M. (1999). Gender electrified: ERP evidence on the syntactic nature of gender processing. Journal of psycholinguistic research, 28, 715–728.

26. Hagoort, P., & Brown, C. M. (2000). ERP effects of listening to speech compared to reading: the P600/SPS to syntactic violations in spoken sentences and rapid serial visual presentation. Neuropsychologia, 38(11), 1531–1549.

27. Hinojosa, J., Martín-Loeches, M., Casado, P., Munoz, F., & Rubia, F. (2003). Similarities and differences between phrase structure and morphosyntactic violations in Spanish: An event-related potentials study. Language and Cognitive Processes, 18(2), 113–142.

28. Iriarte, J., Urrestarazu, E., Valencia, M., Alegre, M., Malanda, A., Viteri, C., & Artieda, J. (2003). Independent component analysis as a tool to eliminate artifacts in EEG: a quantitative study. Journal of clinical neurophysiology, 20(4), 249–257.

29. Jayaseelan, K. A. (2004). 11. The possessor — experiencer dative in Malayalam. In Typological Studies in Language (pp. 227–244). John Benjamins Publishing Company. 10.1075/tsl.60.13jay

30. Kaan, E. (2002). Investigating the effects of distance and number interference in processing subject-verb dependencies: An ERP study. Journal of Psycholinguistic Research, 31, 165–193.

31. Kemmerer, D., Castillo, J. G., Talavage, T., Patterson, S., & Wiley, C. (2008). Neuroanatomical distribution of five semantic components of verbs: Evidence from fMRI. Brain and language, 107(1), 16–43.

32. Kluender, R., & Kutas, M. (1993). Bridging the gap: Evidence from ERPs on the processing of unbounded dependencies. Journal of cognitive neuroscience, 5(2), 196–214.

33. Kretzschmar, F., Bornkessel-Schlesewsky, I., Staub, A., Roehm, D., & Schlesewsky, M. (2011). Prominence Facilitates Ambiguity Resolution: On the Interaction Between Referentiality, Thematic Roles and Word Order in Syntactic Reanalysis. In Studies in Theoretical Psycholinguistics (pp. 239–271). Springer Netherlands. 10.1007/978-94-007-1463-2_11

34. Leinonen, A., Brattico, P., Järvenpää, M., & Krause, C. M. (2008). Event-related potential (ERP) responses to violations of inflectional and derivational rules of Finnish. Brain research, 1218, 181–193.

35. Lenth Russell, V. (2021). emmeans: Estimated Marginal Means, aka Least-Squares Means. R package version 1.6. 0.

36. Mateu, V. (2022). Object wh-questions with psych verbs are easy in child Spanish. Y. Gong, &McKoon, G., & Macfarland, T. (2002). Event templates in the lexical representations of verbs. In Cognitive Psychology (Vol. 45, Issue 1, pp. 1–44). Elsevier BV. 10.1016/s0010-0285(02)00004-x

37. Molinaro, N., Barber, H. A., & Carreiras, M. (2011). Grammatical agreement processing in reading: ERP findings and future directions. cortex, 47(8), 908–930.

38. Münte, T. F., Heinze, H. J., Matzke, M., Wieringa, B. M., & Johannes, S. (1998). Brain potentials and syntactic violations revisited: No evidence for specificity of the syntactic positive shift. Neuropsychologia, 36(3), 217–226.

39. Muraki, E. J., Cortese, F., Protzner, A. B., & Pexman, P. M. (2020). Heterogeneity in abstract verbs: An ERP study. Brain and language, 211, 104863.

40. Muralikrishnan, R., & Idrissi, A. (2021). Salience-weighted agreement feature hierarchy modulates language comprehension. In Cortex (Vol. 141, pp. 168–189). Elsevier BV. 10.1016/j.cortex.2021.03.029

41. Nicenboim, B. (2018). eeguana: A package for manipulating EEG data in R. Computer software Version 0.1.11.9001. *Retrieved from* https://github*. com/bnicenboim/eeguana*. 10.5281/zenodo.2533138

42. Nieuwland, M. S., Martin, A. E., & Carreiras, M. (2013). Event-related brain potential evidence for animacy processing asymmetries during sentence comprehension. In Brain and Language (Vol. 126, Issue 2, pp. 151–158). Elsevier BV. 10.1016/j.bandl.2013.04.005

43. Perani, D., Cappa, S. F., Schnur, T., Tettamanti, M., Collina, S., Rosa, M. M., & Faziol, F. (1999). The neural correlates of verb and noun processing: A PET study. Brain, 122(12), 2337-2344.

44. Pion-Tonachini, L., Kreutz-Delgado, K., & Makeig, S. (2019). ICLabel: An automated electroencephalographic independent component classifier, dataset, and website. NeuroImage, 198, 181–197.

45. Rafi, Z., & Greenland, S. (2020). Semantic and cognitive tools to aid statistical science: replace confidence and significance by compatibility and surprise. In BMC Medical Research Methodology (Vol. 20, Issue 1). Springer Science and Business Media LLC. 10.1186/s12874-020-01105-9

46. Repetto, C., Cipresso, P., & Riva, G. (2015). Virtual action and real action have different impacts on comprehension of concrete verbs. Frontiers in psychology, 6, 176.

47. Roehm, D., Winkler, T., Swaab, T., & Klimesch, W. (2002). The N400 and delta oscillations: Is there a difference?. Journal of Cognitive Neuroscience, 134-135.

48. S Shalu, R. Muralikrishnan, Anna Merin Mathew, Kamal Kumar Choudhary. bioRxiv 2025.03.12.642939; doi: 10.1101/2025.03.12.642939

49. Sabsevitz, D. S., Medler, D. A., Seidenberg, M., & Binder, J. R. (2005). Modulation of the semantic system by word imageability. Neuroimage, 27(1), 188–200.

50. Schad, D. J., Vasishth, S., Hohenstein, S., & Kliegl, R. (2020). How to capitalize on a priori contrasts in linear (mixed) models: A tutorial. In Journal of Memory and Language (Vol. 110, p. 104038). Elsevier BV. 10.1016/j.jml.2019.104038

51. Thomas, S. C. (2014). *Understanding the neurophysiological representation patterns of non-verifiable mental action verbs: an ERP investigation* (Doctoral dissertation, Laurentian University of Sudbury).

52. Thomas, S. C., Brothers, T., Vares, D., & Dickinson, J. (2012, December). Verifiability and Action verb Processing: An ERP Investigation. In CANADIAN JOURNAL OF EXPERIMENTAL PSYCHOLOGY-REVUE CANADIENNE DE PSYCHOLOGIE EXPERIMENTALE (Vol. 66, No. 4, pp. 280-280). 141 LAURIER AVE WEST, STE 702, OTTAWA, ONTARIO K1P 5J3, CANADA: CANADIAN PSYCHOLOGICAL ASSOC.

53. Wasserstein, R. L., Schirm, A. L., & Lazar, N. A. (2019). Moving to a World Beyond “*p* < 0.05.” The American Statistician, 73(sup1), 1–19. 10.1080/00031305.2019.1583913

54. Wilson, M., & Dillon, B. (2022). Alignment between Thematic Roles and Grammatical Functions Facilitates Sentence Processing: Evidence from Experiencer Verbs. In SSRN Electronic Journal. Elsevier BV. 10.2139/ssrn.4235952

